# The kinesin motor KIF1C is a putative transporter of the exon junction complex in neuronal cells

**DOI:** 10.1101/2022.08.24.505074

**Authors:** Maike Nagel, Marvin Noß, Jishu Xu, Nicola Horn, Marius Ueffing, Karsten Boldt, Rebecca Schüle

**Author notes:** **Correspondence:** Rebecca Schüle, Department of Neurodegenerative Diseases, University of Tübingen, Hoppe-Seyler-Str. 3, 72076 Tübingen, Germany.

## Abstract

Neurons critically depend on regulated RNA localization and tight control of spatio-temporal gene expression to maintain their morphological and functional integrity. Mutations in the kinesin motor protein gene *KIF1C* cause Hereditary Spastic Paraplegia, an autosomal recessive disease leading to predominant degeneration of the long axons of central motoneurons. In this study we aimed to gain insight into the molecular function of KIF1C and understand how KIF1C dysfunction contributes to motoneuron degeneration.

We used affinity proteomics in neuronally differentiated neuroblastoma cells (SH-SY5Y) to identify the protein complex associated with KIF1C in neuronal cells; candidate interactions were then validated by immunoprecipitation and mislocalization of putative KIF1C cargoes was studied by immunostainings.

We found KIF1C to interact with all core components of the exon junction complex (EJC); expression of mutant KIF1C in neuronal cells leads to loss of the typical localization distally in neurites. Instead, EJC core components accumulate in the pericentrosomal region, here co-localizing with mutant KIF1C. These findings suggest KIF1C as a neuronal transporter of the EJC. Interestingly, the binding of KIF1C to the EJC is RNA-mediated, as treatment with RNAse prior to immunoprecipitation almost completely abolishes the interaction. Silica-based solid-phase extraction of UV-crosslinked RNA-protein complexes furthermore supports direct interaction of KIF1C with RNA, as recently also demonstrated for kinesin heavy chain. Taken together, our findings are consistent with a model where KIF1C transports mRNA in an EJC-bound and therefore transcriptionally silenced state along neurites, thus providing the missing link between the EJC and mRNA localization in neurons.

## Introduction

Neurons critically rely on precisely regulated spatio-temporal gene expression and protein translation as they are highly polarized, display extreme morphological complexity and require a potent communication between their soma and distal processes (Bentley and Banker 2016). Consequently, localization of mRNA and subsequent local protein synthesis contribute to multiple neuronal functions like synaptogenesis, synapse pruning, axon guidance, axon regeneration, and synaptic plasticity (Sutton and Schuman 2006; Wang et al. 2010; Perry and Fainzilber 2014; Tom Dieck et al. 2014). Axons and dendrites thus display a highly complex transcriptome, with more than 2,000 mRNAs localizing to neurites of hippocampal neurons (Cajigas et al. 2012; Kar et al. 2018).

The exon junction complex (EJC) has been suggested to link transcription, splicing, nonsense-mediated mRNA decay, cytoplasmic mRNA localization and local translation (Woodward et al. 2017). Hereby, the EJC assembles upstream of exon-junctions during the nuclear splicing of pre-mRNA and remains bound to the RNA throughout most its lifecycle until it is removed during the ‘pioneer round of translation’. The eukaryotic Translation Initiation Factor 4A3 (eIF4A3) is the anchor around which the EJC assembles; mago nashi homolog (MAGOH) and RNA-binding protein 8A / Y14 (RBM8A) have been identified as additional core components of the complex (Chan et al. 2004; Ferraiuolo et al. 2004). In mammals, MAGOH protein may be synthesized from either of two homologous genes *MAGOH* and *MAGOHB* (Boehm and Gehring 2016). Metastatic lymph node gene 51 protein (MLN51/CASC3/Barentsz) was proposed as a fourth core component, but recent studies suggest that it is not required for EJC assembly during splicing in living cells and thus is no compulsory component (Mabin et al. 2018; Gerbracht et al. 2020; Schlautmann and Gehring 2020). Additionally, more than 30 proteins were specifically purified with the EJC core and identified as so-called peripheral EJC components (Tange et al. 2005; Merz et al. 2007; Singh et al. 2012).

Little is known about the specific role of the EJC in mammalian neurons. Studies suggest a role in mRNA transport and localization, although most evidence is indirect. In neuronal processes, messenger ribonucleoproteins (mRNPs) containing components of the EJC have been described. These mRNPs constitute functional transport granules which have been detected alongside microtubules (St Johnston 2005; Kiebler and Bassell 2006) and contain translationally silenced mRNAs together with components of the translation machinery. In rat hippocampal and cortical neurons, eIF4A3 showed a somatodendritic localization and associated with other dendritically localized mRNP proteins like Staufen 1 or Fragile X protein. Moreover, eIF4A3 associates with specific dendritic mRNAs (Arc, MAP2, Dendrin) and was necessary to regulate Arc abundance (Giorgi et al. 2007). Further support for a potential link between the EJC and mRNA localization was also obtained in other model systems. In drosophila, the EJC is essential for localization of *oskar* mRNA to the posterior pole of drosophila oocytes (Hachet and Ephrussi 2001; Hachet and Ephrussi 2004; Besse and Ephrussi 2008). In human retinal pigment epithelial (RPE1) cells, EJC components eIF4A3 and RBM8A associate with ninein (NIN) mRNA and are required to maintain the centrosomal localization of NIN transcripts and enable ciliogenesis (Kwon et al. 2021).

The EJC plays important roles in neurodevelopment and deficiency of EJC core components has been shown to lead to defects of embryonic neurogenesis and microcephaly in mice (Bartkowska et al. 2018). Furthermore, mutations in components of the EJC core complex have been linked to two human diseases (*RBM8A* - Thrombocytopenia-absent radius syndrome / #274000; *EIF4A3* - Robin sequence with cleft mandible and limb anomalies / #268305). Moreover, microdeletions of 1q21.1, a locus harboring the *RBM8A* gene, have been associated with autism spectrum disorders (Woodward et al. 2017).

Axonopathies are degenerative disorders primarily characterized by length-dependent degeneration of central or peripheral axonal processes. They comprise Hereditary Spastic Paraplegias (HSP), a large heterogeneous group of genetic degenerative diseases affecting central motor axons in the corticospinal tract and Charcot-Marie-Tooth disease (CMT), peripheral axonopathies with putative sensory as well as motor involvement. The long neuronal axons involved in HSPs and/or CMTs are particularly vulnerable to disturbances of intracellular long-distance transport. Indeed, mutations in multiple members of the kinesin family have been implicated in causing axonopathy phenotypes, including conventional kinesin / kinesin-1 (*KIF5A* – spastic paraplegia type 10 / #604187; *KIF5C* – complex cortical dysplasia with other brain malformations type 2 / #615282; *KLC2* - spastic paraplegia, optic atrophy, and neuropathy / #609541), kinesin-3 (*KIF1A* - hereditary sensory neuropathy type IIC / #614213 and spastic paraplegia type 30 / #610357; *KIF1B* - Charcot-Marie-Tooth disease, axonal, type 2a1 / #118210; *KIF1C* - Spastic ataxia type 2 / #611302), and kinesin-11 (*KIF26B* - autosomal dominant spinocerebellar ataxia and pontocerebellar hypoplasia with arthrogryposis) (Kalantari and Filges 2020). Mutations in the kinesin-3 gene *KIF1C* have been shown to cause autosomal recessive HSP complicated by cerebellar involvement (Caballero Oteyza et al. 2014; Dor et al. 2014; Yucel-Yilmaz et al. 2018; Marchionni et al. 2019; Santos et al. 2022). Mutation types observed in patients with HSP-KIF1C include missense variants in the kinesin-3 motor domain as well as truncating mutations scattered throughout the open reading frame and thus suggest loss of motor function as the relevant disease mechanism. KIF1C is the fastest human cargo transporter (Lipka et al. 2016) and is therefore essential for long distance transport as well as highly dynamic transport requirements.

KIF1C has been connected to multiple cellular functions. Together with dynein it participates in the bidirectional transport of Rab6-positive secretory vesicles along axons and dendrites in rat primary hippocampal neurons (Schlager et al. 2010; Lipka et al. 2016). KIF1C motor activity is regulated by binding to the Rab6 effector BICDR-1 which supports pericentrosomal localization of Rab6-positive vesicles (Schlager et al. 2010). While high levels of BICDR-1 expression in young neurons favor minus-end directed transport of Rab6-positive vesicles, decreasing BICDR-1 levels in maturing neurons shift the balance towards anterograde transport of secretory vesicles required for neurite outgrowth. Accordingly, it has been shown that knockdown of KIF1C in rat primary hippocampal neurons leads to defects of neurite outgrowth (Schlager et al. 2010). Rab6 binding sites at both the C-terminus as well as the N-terminus of KIF1C further mediate a role for KIF1C in maintenance of Golgi morphology which is independent from KIF1C’s motor function; KIF1C depletion consequently leads to fragmentation of the Golgi in HeLa cells (Simpson et al. 2012; Lee et al. 2015). Additionally, KIF1C has been implicated in cell migration. KIF1C-dependent transport of integrins (α5β1-integrins) stabilizes trailing adhesions in human retinal pigment epithelial cells and thus supports directional migration (Theisen et al. 2012). Moreover, KIF1C is involved in formation of podosomes in vascular smooth muscle cells and macrophages (Kopp et al. 2006; Bhuwania et al. 2014; Efimova et al. 2014), a process that has been repeatedly linked to cell migration (Efimova et al. 2014). Furthermore, KIF1C depletion leads to decreased presentation of MHC class II proteins at the cell surface of antigen presenting cells (Franzini et al. 2012). Lastly, KIF1C has recently been implicated in transport of APC-dependent mRNAs. In Hela and human breast cancer MDA-MB-231 cells, KIF1C associates with APC and is required for the active transport of this specific subset of mRNAs (Pichon et al. 2021).

In this study, we aimed at understanding how KIF1C dysfunction leads to neuronal degeneration. Interactome analysis in a neuronal cell model identified interaction of KIF1C with multiple components of the EJC. Hereby, formation of the KIF1C-EJC complex requires binding of RNA. KIF1C mutations lead to mislocalization of EJC components in neuronal cells, suggesting KIF1C as a putative transporter of the EJC in neurons.

## Results

### Affinity proteomics in a neuronal cell model reveals interaction of KIF1C with components of the exon junction complex

The biological role of KIF1C is incompletely understood. To characterize the protein complex interacting with KIF1C in a neuronal cellular background, we therefore performed affinity proteomics in neuronally differentiated SH-SY5Y cells. We overexpressed KIF1C_WT_ and mutant KIF1C_G102A,_ which has been shown to cause hereditary spastic paraplegia (HSP-KIF1C) (Caballero Oteyza et al. 2014), fused to an HA-tag, using expression of an ‘empty’ HA-tag as a control. After affinity purification, KIF1C interactors were subjected to mass spectrometric analysis (Supplementary table 1). 14 proteins were significantly enriched in KIF1C_WT_ expressing cells and 12 out of the 14 proteins were also significantly enriched in KIF1C_G102A_ cells (analysis compared to empty HA-Tag; list of significant interactors in Table 1). Hereby, KIF1C was found to interact with itself; this interaction was to be expected as KIF1C forms dimers (Dorner et al. 1999). 14-3-3 protein theta (YWHAQ), another potential interactor of KIF1C identified in a previous proteome analysis (Wang et al. 2011) was also re-identified in our analysis. Pathway enrichment analysis using WebGestalt revealed that 9 of the significant interactors are associated with the exon junction complex (p= 0.0) (Fig. 1 A). Furthermore, the GO terms RNA binding (p=1,64e^−6^; 12 out of the 14 interactors), mRNA processing (p= 1.4433e^−14^; 11/14) and RNA localization (p= 3.3785e^−8^; 6/14) were enriched. All core components of the EJC, including eIF4A3, RBM8A and MAGOH, were found among the significantly enriched hits, as well as some peripheral components of the EJC like CASC3, Chromatin target of PRMT1 (CHTOP) and all components of the apoptosis- and splicing-associated ASAP and PSAP complexes.

**Table 1:**
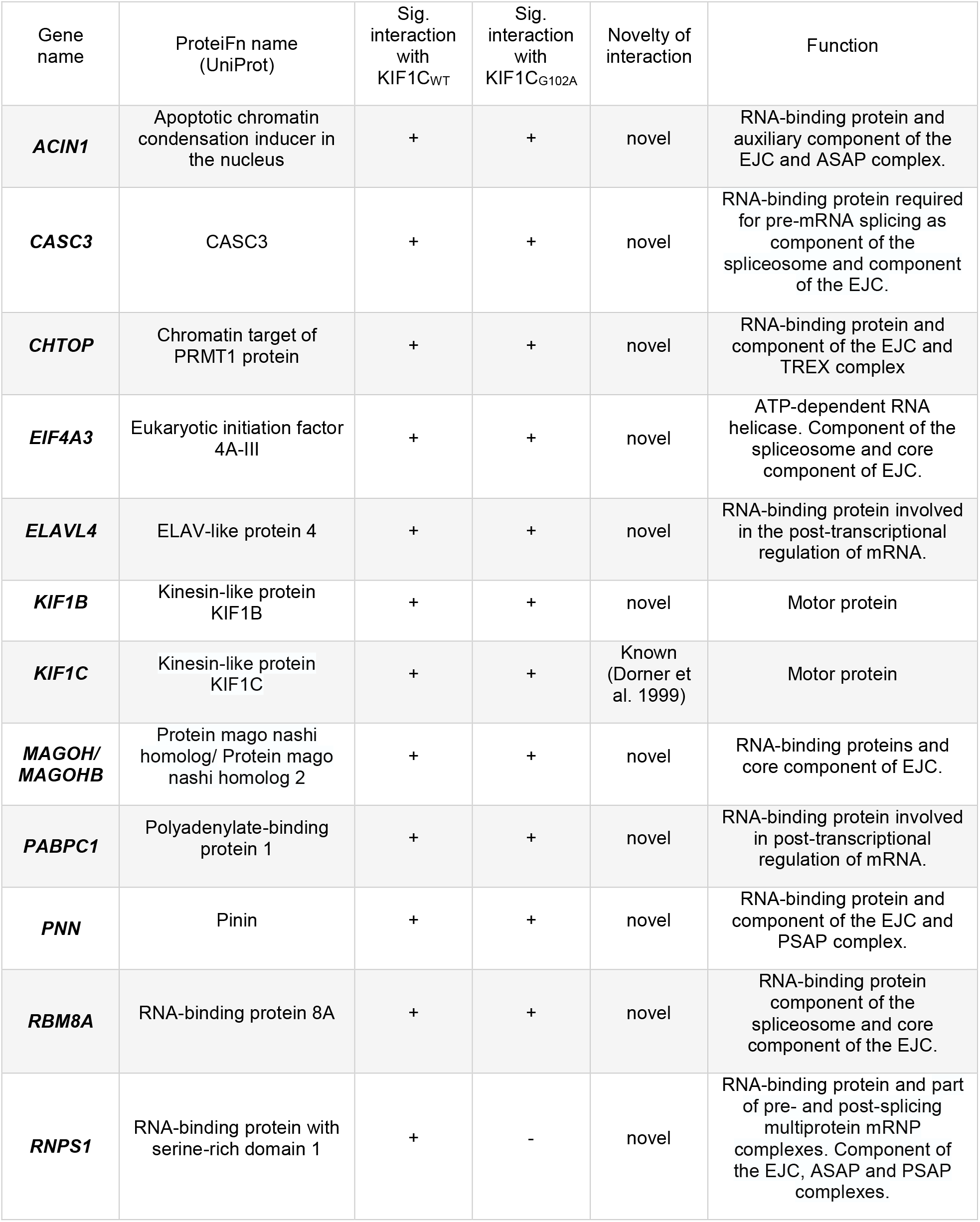
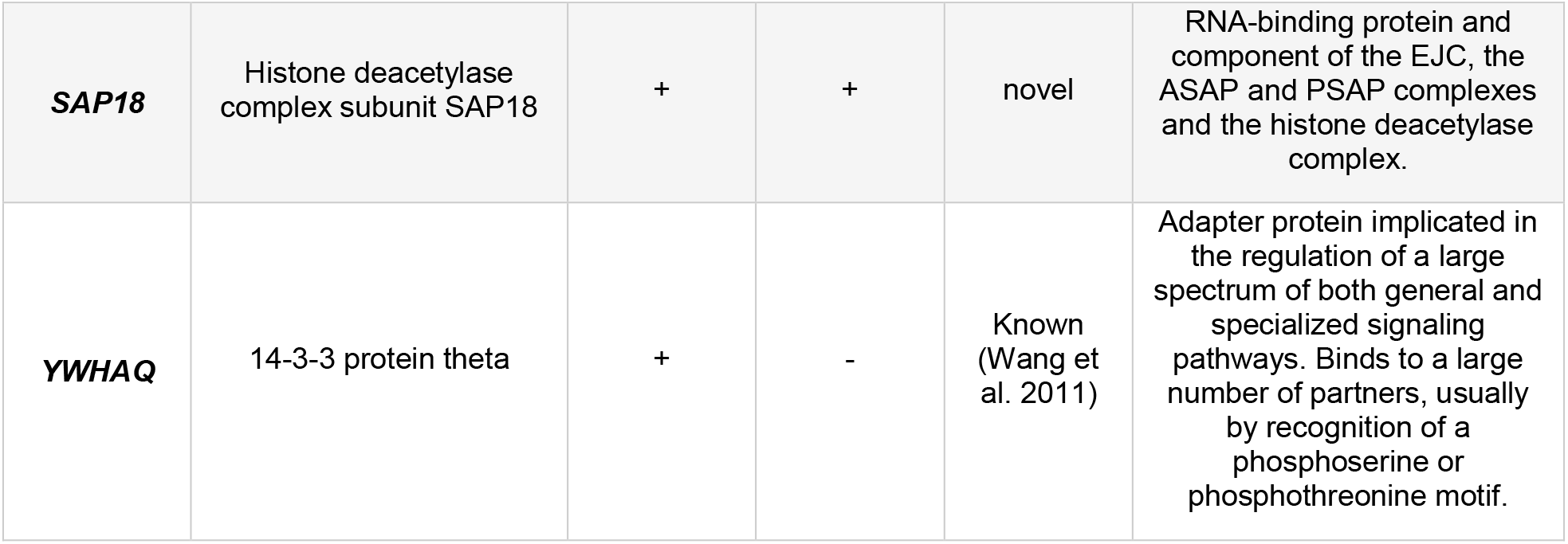
Overview of significant MS hits. Proteins interacting with KIF1C according to MS analysis in differentiated SH-SY5Y cells. Integrators are filtered by significance (p < 0.05 in both Significance A and permutation-based corrected t-test).

**Figure 1:**
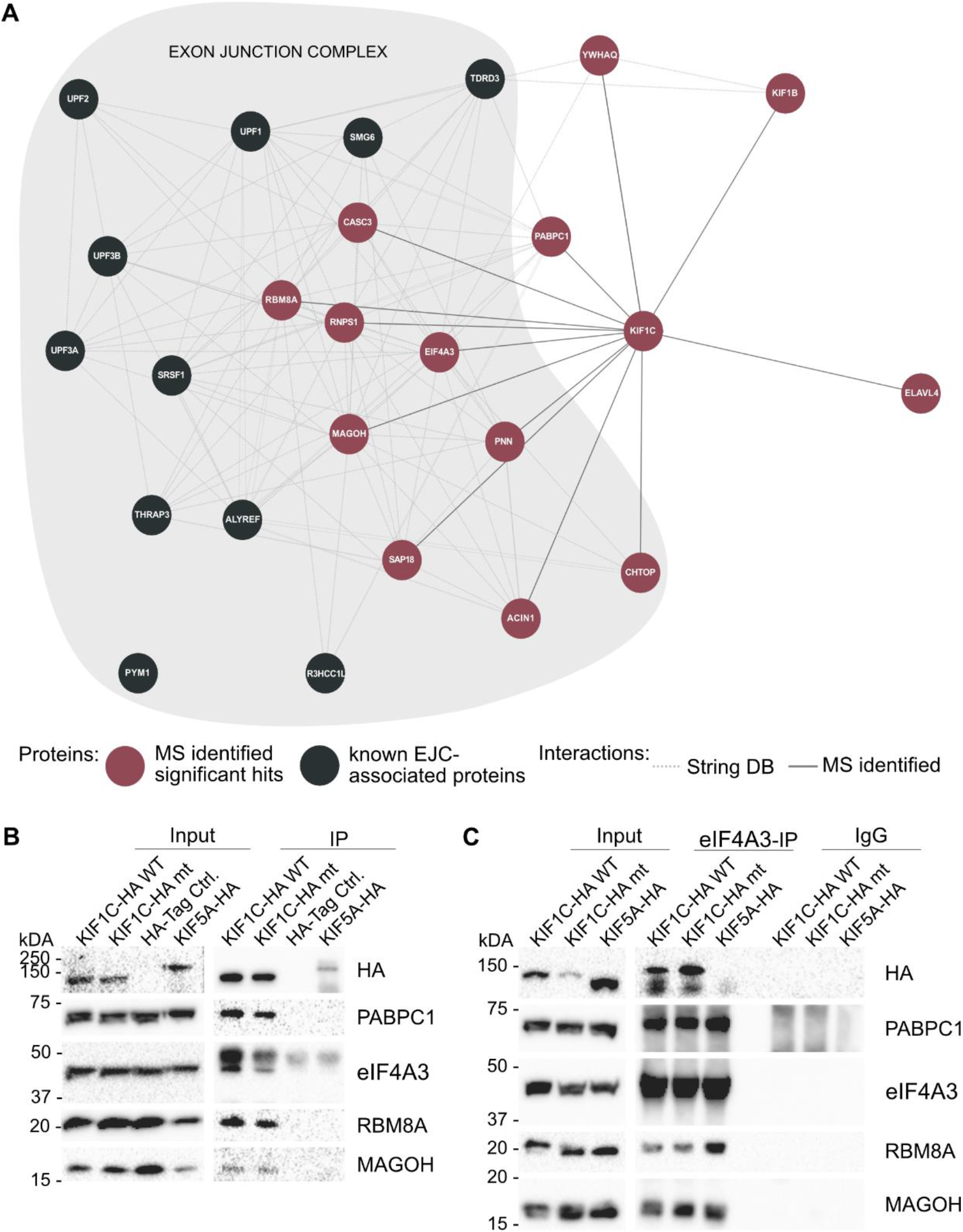
Interaction of exon junction complex components and KIF1C. **A:** Protein-protein-interaction network of KIF1C: KIF1C interactors identified in this study are indicated in red; additional proteins associated with the exon junction complex (according to GO terms) are depicted in grey. **B:** Immunoprecipitation of KIF1C complexes in differentiated SH-SY5Y cells stably overexpressing KIF1C-HA constructs. Interaction with new interaction partners identified by MS was confirmed in the KIF1C IPs; as expected these proteins do not interact with the HA-Tag and wildtyp KIF5A-HA (control). **C:** Co-immunoprecipitation of eIF4A3 in differentiated SH-SY5Y cell lines. KIF1C_WT_ and KIF1C_G102A_ were detected in the Co-IP, while KIF5A-HA (control) was missing. All other tested proteins are known to interact with eIF4A3 and were accordingly identified in all IP samples.

Interestingly, additional peripheral EJC components were also found to be enriched, albeit with slightly less stringent filter criteria (Supplementary table 1). Enrichment of EJC associated proteins (p = 0.0) and RNA-associated GO terms was also confirmed for this extended list (110 proteins total), which additionally included the EJC associated proteins THO complex subunit 4 (ALY/REF), Telomerase-binding protein EST1A (SMG6), Thyroid hormone receptor associated protein 3 (THRAP3) and Serine/arginine-rich splicing factor 1 (SRSF1).

### Immunoprecipitation validates the interaction of KIF1C with PABP, eIF4A3, RBM8A and MAGOH

To validate the putative interaction of KIF1C with the EJC, we performed an immunoprecipitation (IP) with the three obligatory core components of the EJC – eIF4A3, RBM8A and MAGOH – as well as with Polyadenylate-binding protein 1 (PABPC1), an RNA binding protein (RBP) that has been repeatedly shown to interact with components of the EJC (Kashima et al. 2006; Izumi et al. 2010; Singh et al. 2012). All four selected interaction partners identified by mass spectrometry could be confirmed by IP in neuronally differentiated SH-SY5Y cells (Fig. 1 B). None of the tested proteins interacted with the kinesin motor protein KIF5A, mutated in HSP type SPG10, thus confirming that interaction with the EJC is not a class effect but rather appears to be specific for KIF1C. Additionally, no difference could be detected between KIF1C_WT_ and mutant KIF1C_G102A_ (Fig. 1 B) in the IP.

Interaction of KIF1C with eIF4A3 and PABPC1 was furthermore confirmed in a non-neuronal cell model (HEK 293 cells; Supplementary Fig. 1 A).

### Co-immunoprecipitation of core EJC component eIF4A3 confirms interaction

To further confirm the interaction of KIF1C with the EJC, a co-immunoprecipitation (Co-IP) pulling down the endogenously expressed EJC core component eIF4A3 was performed in neuronally differentiated SH-SY5Y cells overexpressing KIF1C variants. Interaction of endogenous eIF4A3 with both wildtype and mutant KIF1C was confirmed (Fig. 1C). Additionally, the known interaction of eIF4A3 with MAGOH and RBM8A could be corroborated. In keeping with the results of the IP, no interaction of eIF4A3 with KIF5A_WT_ was found, again substantiating specificity of the interaction for KIF1C (Fig. 1 C). Subcellular fractionation of differentiated SH-SY5Y cells further confirmed presence of selected EJC core components (eIF4A3, MAGOH) and KIF1C in the cytoplasm. As expected, EJC core components could additionally be detected in the nuclear fraction (Supplementary Fig. 2 A). The presence of eIF4A3 and MAGOH and KIF1C in the cytoplasm was furthermore confirmed in HEK 293 cells and the CO-IP results could additionally be validated in this non-neuronal cell model (Supplementary Fig. 1 C and D).

**Figure 2:**
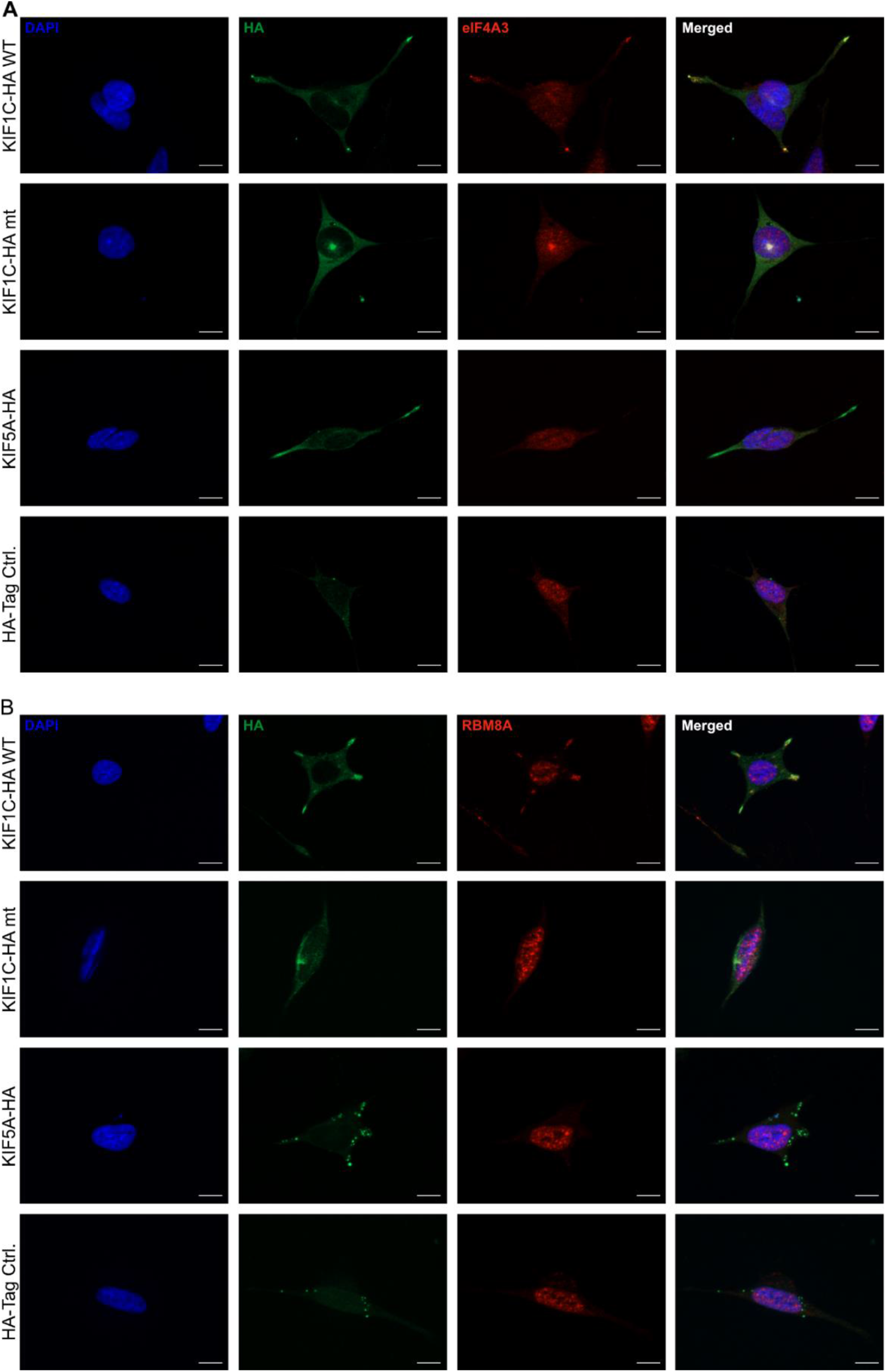
Co-localization of KIF1C and EJC components eIF4A3 and RBM8A. Images show the co-localization of KIF1C_WT_ and the EJC components at the tips of cellular processes. In contrast, KIF1C_G102A_ and the EJC components co-localize in the pericentrosomal region. The scale bars represent 10 μm.

### EJC components mislocalize in neuronal cells overexpressing mutant KIF1C_G102A_

Previous reports have shown that the EJC is involved also in mRNA localization; hereby, EJC components are crucial for the localization of *oscar* mRNA in drosophila oocytes (Hachet and Ephrussi 2001; Hachet and Ephrussi 2004). To investigate whether cellular localization of EJC components depends on KIF1C motility, thus supporting a function of KIF1C in transporting the EJC, we investigated the co-localization of KIF1C and interactors immunocytochemically. Overexpressed KIF1C_WT_ localizes to the tips of cellular protrusion in differentiated SH-SY5Y cells. In contrast, KIF1C_G102A_ is seen as a bright spot next to the nucleus, previously shown to correspond to the pericentrosome (Supplementary Fig. 1 B and Fig. 2 A and B) (Caballero Oteyza et al. 2014). Endogenous eIF4A3 and RBM8A hereby follow the localization pattern of KIF1C: They are found at the tips of cellular protrusions in KIF1C_WT_-overexpressing cells and localize at the centrosome in cells overexpressing KIF1C_G102A_ (Fig. 2 A and B).

### Interaction of KIF1C and the EJC components is RNA-mediated

As the EJC is known to interact with RNA and RNA binding proteins are significantly enriched among the KIF1C interaction partners, we were interested whether the observed interaction of KIF1C with the EJC components is RNA mediated. Thus, we added either RNAse I or RNAse inhibitor to the IP and hypothesized that addition of RNAse will abolish interaction between KIF1C and interaction partners.

RNA depletion during the lysis of differentiated SH-SY5Y cells overexpressing KIF1C resulted in dramatically reduced binding of EJC components and PABPC1 to KIF1C_WT_ and KIF1C_G102A_, whereas RNAse inhibition preserved the interaction (Fig. 3 A). The results could be validated in HEK 293 cells overexpressing KIF1C variants (Supplementary Fig. 1 B).

**Figure 3:**
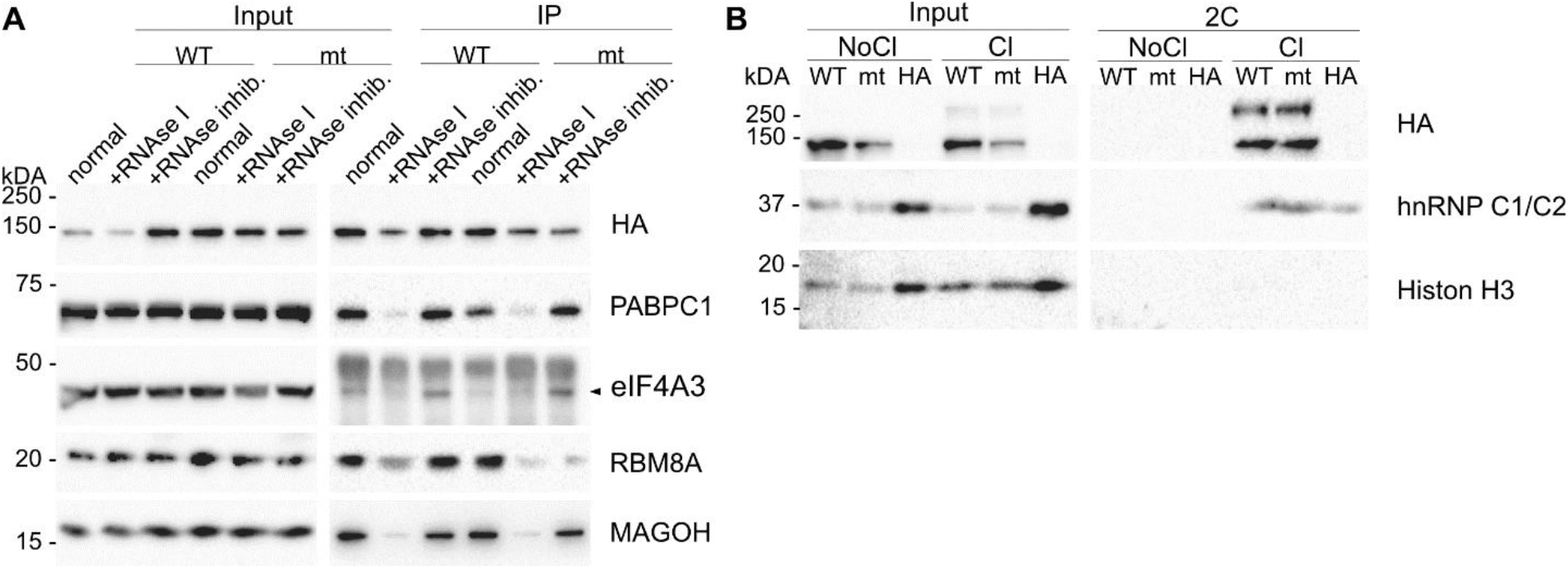
RNA mediated interaction of KIF1C and the EJC. **A:** RNAse IP was performed in differentiated SH-SY5Y cells overexpressing HA-tagged KIF1C_WT_ or KIF1C_G102A_. The proteins interact in a normal IP with KIF1C and the interaction is lost when the samples are treated with RNAse I before the IP, whereas a RNAse inhibition can restore the interaction. **B:** Complex Capture (2C) in HEK 293 cells overexpressing HA-tagged KIF1C_WT_ or KIF1C_G102A_. The input shows signal in crosslinked (Cl) and not crosslinked (NoCl) samples, whereas in the 2C samples only proteins crosslinked to RNA are detected. Histon H3 is a DNA binding protein and used as a control to confirm the specificity of the method for RNA-protein interactions. As a positive control, the known RNA binding protein hnRNP C1/C2 was used. It is detected in the 2C crosslinked samples as well as wildtype and mutant KIF1C.

### KIF1C binds RNA

As the RNAse IP suggested a possible interaction of KIF1C with RNA, we sought independent confirmation via ‘complex capture’ (2C), a method to enrich nucleic-acid bound proteins making them accessible to western blot analysis (Asencio et al. 2018). We therefore overexpressed KIF1C variants in HEK 293 cells and crosslinked proteins and nucleic acids with “zero distance” by UV radiation, thus causing DNA and RNA to covalently bind proteins nearby. After cell lysis, crosslinked protein-RNA complexes were extracted using silica-based solid-phase extraction, whereby we exploited the property of the silica matrix to retain nucleic acids. To favor RNA over DNA ethanol is added to the samples and rigorous washing steps are included.

Using 2C in HEK 293 cells overexpressing KIF1C we were thus able to demonstrate binding of both KIF1C_WT_ and KIF1C_G102A_ to RNA (Fig. 3 B). Due to the property of the 2C protocol – UV crosslinking typically occurs at 5 – 30 Å and thus requires very close nucleic acid / protein contact – this interaction is likely to reflect direct binding (Harris and Christian 2009).

## Discussion

We have here shown for the first time that the motor protein KIF1C associates with components of the EJC in neuronal cells. KIF1C interacts with all three members of the EJC core complex: eIF4A3, MAGOH and RBM8A (validated via IP) as well as CASC3 (identified by MS analysis), a protein stably interacting with EJC components (Degot et al. 2004). The EJC core complex assembles in the nucleus; assembly is initiated by recruitment of eIF4A3 to the spliceosomal protein CWC22. The following interaction of eIF4A3 with the ribose-phosphate backbone of RNA (Andersen et al. 2006; Bono et al. 2006) involves an ATP dependent conformational change in eIF4A3 and interaction to the MAGOH-RBM8A heterodimer (Ballut et al. 2005; Schlautmann and Gehring 2020). The resulting complex is highly stable and – although formation requires presence of RNA – interaction of the core complex proteins is maintained even after RNAse digestion (Singh et al. 2012). Moreover, it has been demonstrated that the complex does not assemble after cell lysis (Singh et al. 2012). It is therefore likely that KIF1C indeed interacts with the EJC core complex rather than individual core complex components. Among the numerous peripheral factors associated with the EJC through its lifecycle, we additionally identified two specific subcomplexes to interact with KIF1C using the most stringent filter settings: the apoptosis- and splicing-associated protein (ASAP) complex, comprising the proteins SAP18, RNPS1 and Acinus (*ACIN1*) and the PSAP complex, in which Acinus is replaced by Pinin (PNN). Although we cannot completely rule out the possibility that these peripheral factors may have associated with the EJC core complex post cell lysis, several arguments support the interpretation that KIF1C indeed interacts with a version of the EJC containing the ASAP and/or PSAP complex: (i) only for those two subcomplexes we identified all components in our screening using stringent filter criteria; (ii) in previous affinity proteomic screens, numerous additional peripheral factors where identified (e.g. Pym1; TREX components CHTOP, DDX39B, UIF, DDX39A or Upfs), when using either eIF4A3 or MAGOH to purify the EJC from HEK 293 cell lysates, indicating that these factors are able to be pulled down from cell lysates. Although some of these components were present in our dataset (see moderate filter set Supplementary table 1), Chromatin target of PRMT1 (CHTOP), another component of the transcription-export (TREX) complex, was the only additional stringently enriched interactor. No other components of the TREX complex, however, were found to be significantly enriched.

ASAP and PSAP are both involved in regulating alternative splicing events but are distinct in their specific functions (Wang et al. 2018). While the two peptides common to both complexes – RNPS1 and SAP18 - have been described to shuttle between the nucleus and the cytoplasm, only nuclear localization has so far been described for Acinus and Pinin (Woodward et al. 2017). However, a growing body of evidence suggests that splicing may also occur in the cytoplasm. In fact, spliceosomal protein and nucleic acid components were identified in dendrites of rat primary neurons and alternative splicing using non-canonical splice sites was demonstrated in dendrites mechanically severed from the soma (Glanzer et al. 2005; Buckley et al. 2014). Moreover, large numbers of intron-retaining transcripts were identified in dendrites (Buckley et al. 2011). It is therefore conceivable that ASAP and PSAP have a role in regulating extranuclear alternative splicing, a function which may be particularly important in neurons.

It is well-established that EJC components are part of transport-competent mRNPs and can be observed dendritically in neurons (Giorgi and Moore 2007; Fritzsche et al. 2013; Wang et al. 2017). mRNPs interact with adaptors and molecular motors to be actively transported throughout the cell to their intended destination (Xing and Bassell 2013; Buchan 2014). Although it is widely assumed that plus-end directed long-range transport of mRNPs along the microtubule cytoskeleton is kinesin dependent, the contribution of individual members of this large family of motor proteins to mRNP transport has not been fully elucidated. We here provide evidence that KIF1C is involved in transport of EJC components along neuronal processes *in vitro* as overexpression of KIF1C carrying a Gly102Ala mutation in the Walker A motif of the nucleotide binding p-loop rendering KIF1C catalytically inactive and thus immobile (Dorner et al. 1998), leads to loss of the peripheral localization of EJC components that can typically be observed when overexpressing wildtype KIF1C in neuronal cells. Previously, studies in several model organisms have supported a contribution of conventional kinesin / kinesin-1 to mRNP transport. Kinesin-1, but also kinesin-2 have been detected in mRNPs isolated from Xenopus oocytes (Messitt et al. 2008; Falley et al. 2009). In hippocampal neurons derived from E16 mouse embryos, kinesin-1 (KIF5A/B/C) transports mRNA-containing granules selectively into dendrites, not axons, whereby binding between kinesin-1 and mRNPs appears to be direct and does not involve kinesin light chain (Kanai et al. 2004). Kinesin-1 (KIF5C) was also reported to transport granules containing shank1 mRNA in dendrites of rat hippocampal neurons, a process that requires presence of the cargo-adapter Staufen1 (Falley et al., 2009). Other kinesin family members implicated in transport of mRNPs include kinesin-2 (FMRP-dependent transport of mRNPs by KIF3C in dendrites of mouse primary hippocampal neurons; (Davidovic et al. 2007)) and kinesin-5 (KIF11; ZBP1-dependent transport of β-actin mRNA in mouse embryo fibroblasts; (Song et al. 2015)). It is therefore possible, that several kinesins in addition to KIF1C contribute to the transport of mRNPs and thus the EJC in neurons. To investigate the individual contribution of individual kinesins to mRNP transport as well as the presence of compensatory mechanisms betweRen different members of the kinesis family, further studies under endogenous conditions as well as *in vivo* would be necessary. It is also not yet clear whether KIF1C specifically transports EJC components or is involved in mRNP transport in general.

Transport of mRNAs via the EJC typically occurs in a translationally repressed state. Cis-acting elements, sequence motifs mostly located in the 3’-UTR of localized RNAs, act as ‘zipcodes’ and direct RNAs to specific cellular locations (Andreassi and Riccio 2009). These sequence motifs interact with trans-acting RNA binding proteins to form transport-competent mRNPs. The EJC core is removed from the mRNA during the initial round of translation (Dostie and Dreyfuss 2002). This translation-dependent disassembly of the EJC is mediated by PYM1 (Partner of Y14 and mago), a protein associated with the 40S ribosome subunit which interacts with the MAGOH-RBM8A heterodimer and thus destabilizes the core of the EJC (Dostie and Dreyfuss 2002). We observe neuritic localization of the EJC in SH-SY5Y cells, this indirectly suggests that mRNAs bound by the KIF1C-EJC complex have not yet undergone the pioneer round of translation. This hypothesis is further supported by studies in differentiated SH-SY5Y cells by Wang et al. Here, the EJC core components eIF4A3 and RBM8A were both found in CBP80-positive neuronally localized mRNP granules (Wang et al. 2017). The nuclear cap binding heterodimer complex CBP80-CBP20 (CBC) and the EJC typically associate with newly synthesized mRNA; during the first round of translation, the EJC is removed, and CBC replaced by eIF4E. Co-occurrence of CBP80 and EJC components in localized mRNP granules was therefore interpreted as indication that some localized mRNAs have not yet undergone their very first translation (Wang et al. 2017). Interestingly, we observed that the interaction between KIF1C and the EJC complex is RNA-mediated as RNA digestion prior to the IP dramatically lowered the affinity of KIF1C to EJC core components. This observation cannot be explained by disassembly of the EJC core complex, as the EJC core complex – once formed – is stable even after RNAse I treatment (Singh et al. 2012). Moreover, the interaction between KIF1C and the RNA-binding protein PABPC1 is equally RNA-dependent. These findings allow two possible interpretations of the nature of the KIF1C-EJC interaction: i) Individual mRNAs might indeed mediate interaction between KIF1C and EJC complex components or PABPC1; this model would not require protein-protein-interactions between KIF1C, EJC components and/or PABPC1. The interaction between KIF1C and mRNA might hereby be direct or mediated via a yet unknown adapter protein (see below). We did, however, not identify significant interaction between KIF1C and any known cargo adaptors (e.g. kinesin light chain 2 (KLC2), Kinesin-associated protein 3 (KIFAP3) and others). ii) Multiple mRNAs, EJC components and other RBPs including PABPC1 might form a macromolecular mRNP which is destabilized upon digestion of RNA and which associates with KIF1C.

Our findings add support to previous evidence from several high-throughput studies (Castello et al. 2012; Dietzel et al. 2012) suggesting direct binding between KIF1C and mRNA as we have demonstrated that KIF1C binds RNA at ‘zero distance’ using UV crosslinking followed by silica-based solid-phase extraction (2C). Interestingly, kinesin heavy chain was recently also shown to directly bind to RNA via an alternative cargo binding domain with high affinity and that this interaction is stabilized, but not mediated by tropomyosin 1 in D. melanogaster (Dimitrova-Paternoga et al. 2021). Direct RNA-binding might therefore be a mechanism of cargo interaction common to several kinesin motors.

In conclusion, our data suggests that KIF1C is involved in transporting the EJC core complex into neurites in a neuronal cell model and that binding between KIF1C and EJC components requires presence of RNA. Preliminary evidence further supports direct binding of KIF1C to RNA. Our findings thus support a model where mRNA is transported along neurites in a transcriptionally silenced state bound to the EJC, thus providing the missing link between the EJC and mRNA localization in neurons.

## Materials and Methods

### Cell culture

Stable cell lines were cultured under standard conditions in Dulbecco’s modified eagle medium (DMEM) high glucose (Sigma-Aldrich), supplemented with 10 % fetal bovine serum (FBS) (Thermo Fisher Scientific) and 1 % neomycin (Sigma-Aldrich) for HEK 293 cells, and in DMEM/F12 (Sigma-Aldrich) with 15 % FBS and 1 % neomycin for SH-SY5Ys. Cells were trypsinised when confluent and plated according to the required cell number. To generate cell lines stably expressing KIF1C, we expressed plasmids containing the HA-tagged wildtype (WT) or mutant (mt) (c.305G>C; p.G102A (Caballero Oteyza et al. 2014) KIF1C open reading frame in a pSF-CMV-UB-NEO/G418 ASCI (Sigma-Aldrich) backbone and cultured cells under neomycin selection conditions. Control cell lines were generated with a plasmid expressing WT HA-tagged KIF5A, a gene mutated in Hereditary Spastic Paraplegia Type SPG10, and a plasmid only expressing the HA-Tag.

To induce neuronal differentiation of SH-SY5Y cells, the medium of 60 % confluent cells was switched to DMEM/F12 with reduced serum concentration (5 % FBS), 1 % neomycin and addition of 10 μM retinoic acid (RA) (Stemcell Technologies). At day 6 the cells were washed twice with warm PBS (Sigma-Aldrich) and the media was changed to serum-free DMEM/F12 with 1 % neomycin and 25 ng/ml BDNF (Peprotech). The cells were used from day 12 to day 18 after differentiation.

### Mass spectrometry

The IP samples were precipitated with chloroform and methanol followed by trypsin digestion as described before (Gloeckner et al. 2009). LC-MS/MS analysis was performed on Ultimate3000 nanoRSLC systems (Thermo Scientific) coupled to an Orbitrap Fusion Tribrid mass spectrometer (Thermo Scientific) by a nano spray ion source. Tryptic peptide mixtures were injected automatically and loaded at a flow rate of 30 μl/min in 0.1 % trifluoroacetic acid in HPLC-grade water onto a nano trap column (300 μm i.d. × 5 mm Pre collumn, packed with Acclaim PepMap100 C18, 5 μm, 100 Å; Thermo Scientific). After 3 minutes, peptides were eluted and separated on the analytical column (75 μm i.d. × 25 cm, Acclaim PepMap RSLC C18, 2 μm, 100 Å; Thermo Scientific) by a linear gradient from 2 % to 30 % of buffer B (80 % acetonitrile and 0.08 % formic acid in HPLC-grade water) in buffer A (2 % acetonitrile and 0.1 % formic acid in HPLC-grade water) at a flow rate of 300 nl/min over 92 minutes. Remaining peptides were eluted by a short gradient from 30 % to 95 % buffer B in 5 minutes. Analysis of the eluted peptides was done on an LTQ Fusion mass spectrometer. From the high resolution MS pre-scan with a mass range of 335 to 1500, the most intense peptide ions were selected for fragment analysis in the ion trap by using a high speed method if they were at least doubly charged. The normalized collision energy for HCD was set to a value of 27 and the resulting fragments were detected with a resolution of 120,000. The lock mass option was activated; the background signal with a mass of 445.12003 was used as lock mass (Olsen et al. 2005). Every ion selected for fragmentation was excluded for 20 seconds by dynamic exclusion. MS/MS data were analyzed using the MaxQuant software (version 1.6.1.0) (Cox and Mann 2008; Cox et al. 2009). As a digesting enzyme, Trypsin/P was selected with maximal 2 missed cleavages. Cysteine carbamidomethylation was set for fixed modifications, and oxidation of methionine and N-terminal acetylation were specified as variable modifications. The data were analyzed by label-free quantification with the minimum ratio count of 3. The first search peptide tolerance was set to 20, the main search peptide tolerance to 4.5 ppm and the re-quantify option was selected. For peptide and protein identification the human subset of the SwissProt database (release 2014_04) was used and contaminants were detected using the MaxQuant contaminant search. A minimum peptide number of 2 and a minimum length of 7 amino acids was tolerated. Unique and razor peptides were used for quantification. The match between run option was enabled with a match time window of 0.7 min and an alignment time window of 20 min. For each sample, 5 biological replicates were analyzed. The statistical analysis including ratio, t-test and significance A calculation was done using the Perseus software (version 1.5.5.3, (Tyanova et al. 2016)). Proteins that were quantified in at least 3 of 5 experiments and met the criteria for significant enrichment (Significance A and permutation-based corrected t-test p<0.05) were considered as potential interactors. The moderate filter set included only Significance A p<0.05. The hits where then analyzed with WebGestalt (Liao et al. 2019) using gene ontology (GO) terms for cellular compartments, molecular and biological function. For the figure, String DB was used to depict the network of the detected 14 significant MS hits together with proteins listed in the GO term exon junction complex.

### Protein isolation

Cells were washed twice with PBS and scraped in the according lysis buffer containing 1x cOmplete protease inhibitor cocktail (PI) (Roche) on ice. Lysates were kept on ice for 30 min vortexing every 5 min. Cell debris were pelleted at 3000-21000 x g at 4°C for 15-30 min. The Pierce™ BCA Protein Assay Kit (Thermo Scientific) was used to determine protein concentrations.

### Western Blotting

The samples were prepared with NuPAGE™ LDS-Sample Buffer (4x) and NuPAGE™ Sample Reducing Agent (10x) (Thermo Scientific) and boiled at 95°C for 10 min. Then the samples were separated using NuPAGE™ MOPS SDS-Running Buffer (Thermo Scientific) and 10 % acrylamide gels and transferred to a 0.45 μm Nitrocellulose membrane (Merck Millipore). After blocking with 5 % milk in TBS-T, the membranes were incubated overnight at 4° C on a rotator with primary antibody in Western Blocking Reagent (Roche), if not stated otherwise. The membranes were washed 3 times with TBS-T for 5 min and secondary antibodies were added in 5 % milk in TBS-T for an hour at room temperature (RT). For detection, Immobilon Western Chemiluminescent HRP Substrate (Merck Millipore) and ChemiDoc XRS+ Imaging System (Biorad) were used.

### Antibodies

Anti-Mago nashi homolog (ab186431 Abcam, rabbit 1:10000); EIF4A3 (67740-1-Ig Proteintech; mouse 1:2000 in milk), HA (#901513 Biolegend; mouse 1:1000), Histone H3 (ab21054 Abcam; HRP conjugated rabbit 1:1000), hnRNP C1/C2 (GTX10294 Biozol, mouse 1:1000), Peroxidase AffiniPure Goat Anti-Mouse/Rabbit IgG (H+L) (111-035-003 Jackson ImmunoResearch; 1:10000), PABPC1 (10970-1-AP Proteintech, rabbit 1:1000 in milk), PABP(10E10) (NB120-6125 Novus Biologicals, mouse 1:1000), RBM8A (ab229573 Abcam, rabbit 1:1000)

### Immunoprecipitation and co-immunoprecipitation

For the immunoprecipitation (IP) of KIF1C and its bound interactors, Pierce™ Anti-HA Magnetic Beads (Thermo Scientific) were used following the manufacturer’s protocol with slight adaptations. After cell lysis (lysis buffer (Nonident P40 (v/v 0.5 %; Roche) in TBS with PI)) proteins were isolated and equal amounts of proteins were incubated with previously washed 25 μl beads in 2-4 ml buffer for 60-90 min at 4°C on a rotator. After 3 washing steps the samples were eluted from the beads via boiling elution. The beads were resuspended in NuPAGE™ LDS-Sample Buffer (4x) and Sample Reducing Agent (10x) and incubated for 5 min at 70°C and 1100 rpm using a ThermoMixer comfort (Eppendorf).

To perform the Co-IP, the Pierce™ Classic Magnetic IP/Co-IP Kit (Thermo Scientific), with EIF4A3 (17504-1-AP Proteintech, rabbit) antibody and a Normal Rabbit IgG (#2729 Cell Signaling) control was used. Cell lysis was performed as described for the HA-IP with the lysis buffer and PI provided in the kit. Equal amounts of protein were incubated with 5 μg of antibody or IgG control at 4°C overnight on a rotator. The IP was performed following the manufacturer’s protocol. 0.25 mg beads were added to each sample and incubated for 2 hours at RT. After the washing steps, samples were eluted via boiling using NuPAGE™ LDS-Sample Buffer (4x), Sample Reducing Agent (10x) and an incubation for 5 min at 95°C and 1100 rpm.

### Immunocytochemistry and imaging

Cells were seeded onto coverslips for subsequent immunocytochemistry experiments. To fix the cells they were washed once with warm PBS and incubated for 20 min at 37°C with 4 % paraformaldehyde (Merck Millipore). After washing, cells were blocked and permeabilized with 5 % normal goat serum (Sigma-Aldrich) in PBS with 0.1 % Triton X-100 (Carl Roth). Primary antibodies were incubated overnight at 4°C and Alexa Fluor-conjugated secondary antibodies (Invitrogen) were incubated for 1 hour at RT. After counterstaining the nuclei with DAPI (Thermo Scientific) the coverslips were mounted with Dako mounting solution (Agilent Dako) onto microscopy. Four to five random fields per cell line and coverslip were imaged using the Leica SP8 confocal microscope (Leica Camera) or the Zeiss Apoptome 2 (Zeiss); hereby, exposure times were kept constant.

### Antibodies

ß-III-Tubulin (T8660 Sigma-Aldrich; mouse 1:1000), Anti-Mago nashi homolog 2 (ab186431 Abcam, rabbit 1:250), EIF4A3 (17504-1-AP Proteintech, rabbit 1:400), HA (#901513 Biolegend; mouse 1:500), RBM8A (ab229573 Abcam, rabbit 1:250)

### RNase IP

The HA-IP described above was combined with a RNAse treatment or RNAse inhibition, respectively. Hereby, SUPERase•In™ RNase Inhibitor (Thermo Scientific) was added to the lysis buffer and vortexing was avoided during cell lysis. The treatment with Ambion™ RNAseI (Thermo Scientific) was performed after lysis at 30°C for 30 min. Subsequently, the HA-IP protocol was followed.

### Complex Capture (2C)

Complex Capture (2C) was performed as described in Asencio et al. (2018) with modifications to confirm interaction of KIF1C with RNA. HEK 293 cells were irradiated with 150 mJ/cm^2^ of UV light at 254 nm in a Crosslinker (Vilber Lourmat BLX-254). Cells were harvested by scraping in PBS, centrifuged at 300 x g for 5 min and the pellets resuspended in lysis buffer (50 mM Tris–HCl (pH 7,4), 100 mM NaCl, 1 % Igepal CA-630, 0,1 % SDS, 0,5 % sodium deoxycholate) supplemented with PI, SUPERase•In™ RNase Inhibitor and DNAseI (Thermo Scientific). Cells were lysed on ice for 30 min and centrifuged for 10 min at max g at 4°C. Then the nucleic-acid bound protein isolation was performed using buffers from the Quick-RNA Miniprep Kit (Zymo Research) and the Zymo-Spin IIICG Column (Zymo Research). After the elution with DNase/RNase free water for 2 min at RT the sample volumes were reduced using a SpeedVac (Thermo Scientific). The RNA concentration was measured using NanoDrop2000 (Thermo Scientific) and the samples were treated with Ambion™ RNAseI for 30 min at 30°C, before preparing them for the analysis via Western Blotting.

### Fractionation

Fractionation was used to analyze different cell fractions of HEK 293 and neuronally differentiated SH-SY5Y cells. Therefore, a protocol based on REAP (Suzuki et al. 2010) was used. After harvesting the cells, the pellet was triturated 10 times with the lysis buffer and an aliquot was kept as the whole cell sample. The rest of the sample was pop-spinned (centrifuge at 10000 x g for 10 sec) and the supernatant was transferred to a new tube and kept on ice as the cytoplasmic fraction. The remaining pellet was washed in lysis buffer and pop-spinned again. After withdrawal of the supernatant the pellet was resuspended in lysis buffer and kept as the nuclear fraction. All samples were treated in an ultrasonic-bath (Bandelin) for 3 minutes before further use.

## Acknowledgements

Research reported in this publication was supported by the Bundesministerium für Bildung und Forschung (BMBF) through funding for the TreatHSP network (grant 01GM1905 to RS), the National Institute of Neurological Disorders and Stroke (NINDS) and the National Institutes of Health (NIH) under Award Number R01NS072248 (RS), the Deutsche Forschungsgemeinschaft (DFG) as part of the PROSPAX consortium under the frame of EJP RD (grant 441409627 to RS), the European Joint Program on Rare Diseases under the EJP RD COFUND action (grant 825575 to RS) and the Horizon 2020 research and innovation program (grant 779257 ‘Solve-RD’ to RS) and the German Center for Neurodegenerative Diseases (DZNE; grant to RS). RS is a member of the European Reference Network for Rare Neurological Diseases - Project ID 739510.

We thank the group of Matthias Hentze, in particular Ina Huppertz and Claudio Asencio Salcedo, for their help with the 2C technique and all questions regarding RNA-binding proteins.

## References

Andersen CB, Ballut L, Johansen JS, Chamieh H, Nielsen KH, Oliveira CL, Pedersen JS, Séraphin B, Le Hir H, Andersen GR. 2006. Structure of the exon junction core complex with a trapped DEAD-box ATPase bound to RNA. Science 313: 1968–1972.

Andreassi C, Riccio A. 2009. To localize or not to localize: mRNA fate is in 3’UTR ends. Trends Cell Biol 19: 465–474.

Asencio C, Chatterjee A, Hentze MW. 2018. Silica-based solid-phase extraction of cross-linked nucleic acid-bound proteins. Life Sci Alliance 1: e201800088.

Ballut L, Marchadier B, Baguet A, Tomasetto C, Séraphin B, Le Hir H. 2005. The exon junction core complex is locked onto RNA by inhibition of eIF4AIII ATPase activity. Nat Struct Mol Biol 12: 861–869.

Bartkowska K, Tepper B, Turlejski K, Djavadian RL. 2018. Roles of the exon junction complex components in the central nervous system: a mini review. Rev Neurosci 29: 817–824.

Bentley M, Banker G. 2016. The cellular mechanisms that maintain neuronal polarity. Nat Rev Neurosci 17: 611–622.

Besse F, Ephrussi A. 2008. Translational control of localized mRNAs: restricting protein synthesis in space and time. Nat Rev Mol Cell Biol 9: 971–980.

Bhuwania R, Castro-Castro A, Linder S. 2014. Microtubule acetylation regulates dynamics of KIF1C-powered vesicles and contact of microtubule plus ends with podosomes. Eur J Cell Biol 93: 424–437.

Boehm V, Gehring NH. 2016. Exon Junction Complexes: Supervising the Gene Expression Assembly Line. Trends Genet 32: 724–735.

Bono F, Ebert J, Lorentzen E, Conti E. 2006. The crystal structure of the exon junction complex reveals how it maintains a stable grip on mRNA. Cell 126: 713–725.

Buchan JR. 2014. mRNP granules. Assembly, function, and connections with disease. RNA Biol 11: 1019–1030.

Buckley PT, Khaladkar M, Kim J, Eberwine J. 2014. Cytoplasmic intron retention, function, splicing, and the sentinel RNA hypothesis. Wiley Interdiscip Rev RNA 5: 223–230.

Buckley PT, Lee MT, Sul JY, Miyashiro KY, Bell TJ, Fisher SA, Kim J, Eberwine J. 2011. Cytoplasmic intron sequence-retaining transcripts can be dendritically targeted via ID element retrotransposons. Neuron 69: 877–884.

Caballero Oteyza A, Battaloglu E, Ocek L, Lindig T, Reichbauer J, Rebelo AP, Gonzalez MA, Zorlu Y, Ozes B, Timmann D et al. 2014. Motor protein mutations cause a new form of hereditary spastic paraplegia. Neurology 82: 2007–2016.

Cajigas IJ, Tushev G, Will TJ, tom Dieck S, Fuerst N, Schuman EM. 2012. The local transcriptome in the synaptic neuropil revealed by deep sequencing and high-resolution imaging. Neuron 74: 453-466.

Castello A, Fischer B, Eichelbaum K, Horos R, Beckmann BM, Strein C, Davey NE, Humphreys DT, Preiss T, Steinmetz LM et al. 2012. Insights into RNA biology from an atlas of mammalian mRNA-binding proteins. Cell 149: 1393–1406.

Chan CC, Dostie J, Diem MD, Feng W, Mann M, Rappsilber J, Dreyfuss G. 2004. eIF4A3 is a novel component of the exon junction complex. RNA 10: 200–209.

Cox J, Mann M. 2008. MaxQuant enables high peptide identification rates, individualized p.p.b.-range mass accuracies and proteome-wide protein quantification. Nat Biotechnol 26: 1367–1372.

Cox J, Matic I, Hilger M, Nagaraj N, Selbach M, Olsen JV, Mann M. 2009. A practical guide to the MaxQuant computational platform for SILAC-based quantitative proteomics. Nat Protoc 4: 698–705.

Davidovic L, Jaglin XH, Lepagnol-Bestel AM, Tremblay S, Simonneau M, Bardoni B, Khandjian EW. 2007. The fragile X mental retardation protein is a molecular adaptor between the neurospecific KIF3C kinesin and dendritic RNA granules. Hum Mol Genet 16: 3047–3058.

Degot S, Le Hir H, Alpy F, Kedinger V, Stoll I, Wendling C, Seraphin B, Rio MC, Tomasetto C. 2004. Association of the breast cancer protein MLN51 with the exon junction complex via its speckle localizer and RNA binding module. J Biol Chem 279: 33702–33715.

Dietzel M, Baltzer PA, Dietzel A, Zoubi R, Groschel T, Burmeister HP, Bogdan M, Kaiser WA. 2012. Artificial Neural Networks for differential diagnosis of breast lesions in MR-Mammography: a systematic approach addressing the influence of network architecture on diagnostic performance using a large clinical database. Eur J Radiol 81: 1508–1513.

Dimitrova-Paternoga L, Jagtap PKA, Cyrklaff A, Vaishali Lapouge K, Sehr P, Perez K, Heber S, Low C, Hennig J et al. 2021. Molecular basis of mRNA transport by a kinesin-1-atypical tropomyosin complex. Genes Dev 35: 976–991.

Dor T, Cinnamon Y, Raymond L, Shaag A, Bouslam N, Bouhouche A, Gaussen M, Meyer V, Durr A, Brice A et al. 2014. KIF1C mutations in two families with hereditary spastic paraparesis and cerebellar dysfunction. J Med Genet 51: 137–142.

Dorner C, Ciossek T, Muller S, Moller PH, Ullrich A, Lammers R. 1998. Characterization of KIF1C, a new kinesin-like protein involved in vesicle transport from the Golgi apparatus to the endoplasmic reticulum. J Biol Chem 273: 20267–20275.

Dorner C, Ullrich A, Haring HU, Lammers R. 1999. The kinesin-like motor protein KIF1C occurs in intact cells as a dimer and associates with proteins of the 14-3-3 family. J Biol Chem 274: 33654–33660.

Dostie J, Dreyfuss G. 2002. Translation is required to remove Y14 from mRNAs in the cytoplasm. Curr Biol 12: 1060–1067.

Efimova N, Grimaldi A, Bachmann A, Frye K, Zhu X, Feoktistov A, Straube A, Kaverina I. 2014. Podosome-regulating kinesin KIF1C translocates to the cell periphery in a CLASP-dependent manner. J Cell Sci 127: 5179–5188.

Falley K, Schütt J, Iglauer P, Menke K, Maas C, Kneussel M, Kindler S, Wouters FS, Richter D, Kreienkamp HJ. 2009. Shank1 mRNA: dendritic transport by kinesin and translational control by the 5’untranslated region. Traffic 10: 844–857.

Ferraiuolo MA, Lee CS, Ler LW, Hsu JL, Costa-Mattioli M, Luo MJ, Reed R, Sonenberg N. 2004. A nuclear translation-like factor eIF4AIII is recruited to the mRNA during splicing and functions in nonsense-mediated decay. Proc Natl Acad Sci U S A 101: 4118–4123.

Franzini L, Ardigo D, Valtuena S, Pellegrini N, Del Rio D, Bianchi MA, Scazzina F, Piatti PM, Brighenti F, Zavaroni I. 2012. Food selection based on high total antioxidant capacity improves endothelial function in a low cardiovascular risk population. Nutr Metab Cardiovasc Dis 22: 50–57.

Fritzsche R, Karra D, Bennett KL, Ang FY, Heraud-Farlow JE, Tolino M, Doyle M, Bauer KE, Thomas S, Planyavsky M et al. 2013. Interactome of two diverse RNA granules links mRNA localization to translational repression in neurons. Cell Rep 5: 1749–1762.

Gerbracht JV, Boehm V, Britto-Borges T, Kallabis S, Wiederstein JL, Ciriello S, Aschemeier DU, Krüger M, Frese CK, Altmüller J et al. 2020. CASC3 promotes transcriptome-wide activation of nonsense-mediated decay by the exon junction complex. Nucleic Acids Res 48: 8626–8644.

Giorgi C, Moore MJ. 2007. The nuclear nurture and cytoplasmic nature of localized mRNPs. Semin Cell Dev Biol 18: 186–193.

Giorgi C, Yeo GW, Stone ME, Katz DB, Burge C, Turrigiano G, Moore MJ. 2007. The EJC factor eIF4AIII modulates synaptic strength and neuronal protein expression. Cell 130: 179–191.

Glanzer J, Miyashiro KY, Sul JY, Barrett L, Belt B, Haydon P, Eberwine J. 2005. RNA splicing capability of live neuronal dendrites. Proc Natl Acad Sci U S A 102: 16859–16864.

Gloeckner CJ, Boldt K, Ueffing M. 2009. Strep/FLAG tandem affinity purification (SF-TAP) to study protein interactions. Curr Protoc Protein Sci Chapter 19: Unit19.20.

Hachet O, Ephrussi A. 2001. Drosophila Y14 shuttles to the posterior of the oocyte and is required for oskar mRNA transport. Curr Biol 11: 1666–1674.

Hachet O, Ephrussi A. 2004. Splicing of oskar RNA in the nucleus is coupled to its cytoplasmic localization. Nature 428: 959–963.

Harris ME, Christian EL. 2009. RNA crosslinking methods. Methods Enzymol 468: 127–146.

Izumi N, Yamashita A, Iwamatsu A, Kurata R, Nakamura H, Saari B, Hirano H, Anderson P, Ohno S. 2010. AAA+ proteins RUVBL1 and RUVBL2 coordinate PIKK activity and function in nonsense-mediated mRNA decay. Sci Signal 3: ra27.

Kalantari S, Filges I. 2020. ‘Kinesinopathies’: emerging role of the kinesin family member genes in birth defects. J Med Genet 57: 797–807.

Kanai Y, Dohmae N, Hirokawa N. 2004. Kinesin transports RNA: isolation and characterization of an RNA-transporting granule. Neuron 43: 513–525.

Kar AN, Lee SJ, Twiss JL. 2018. Expanding Axonal Transcriptome Brings New Functions for Axonally Synthesized Proteins in Health and Disease. Neuroscientist 24: 111–129.

Kashima I, Yamashita A, Izumi N, Kataoka N, Morishita R, Hoshino S, Ohno M, Dreyfuss G, Ohno S. 2006. Binding of a novel SMG-1-Upf1-eRF1-eRF3 complex (SURF) to the exon junction complex triggers Upf1 phosphorylation and nonsense-mediated mRNA decay. Genes Dev 20: 355–367.

Kiebler MA, Bassell GJ. 2006. Neuronal RNA granules: movers and makers. Neuron 51: 685–690.

Kopp P, Lammers R, Aepfelbacher M, Woehlke G, Rudel T, Machuy N, Steffen W, Linder S. 2006. The kinesin KIF1C and microtubule plus ends regulate podosome dynamics in macrophages. Mol Biol Cell 17: 2811–2823.

Kwon OS, Mishra R, Safieddine A, Coleno E, Alasseur Q, Faucourt M, Barbosa I, Bertrand E, Spassky N, Le Hir H. 2021. Exon junction complex dependent mRNA localization is linked to centrosome organization during ciliogenesis. Nat Commun 12: 1351.

Lee PL, Ohlson MB, Pfeffer SR. 2015. Rab6 regulation of the kinesin family KIF1C motor domain contributes to Golgi tethering. Elife 4.

Liao Y, Wang J, Jaehnig EJ, Shi Z, Zhang B. 2019. WebGestalt 2019: gene set analysis toolkit with revamped UIs and APIs. Nucleic Acids Res 47: W199–W205.

Lipka J, Kapitein LC, Jaworski J, Hoogenraad CC. 2016. Microtubule-binding protein doublecortin-like kinase 1 (DCLK1) guides kinesin-3-mediated cargo transport to dendrites. EMBO J 35: 302–318.

Mabin JW, Woodward LA, Patton RD, Yi Z, Jia M, Wysocki VH, Bundschuh R, Singh G. 2018. The Exon Junction Complex Undergoes a Compositional Switch that Alters mRNP Structure and Nonsense-Mediated mRNA Decay Activity. Cell Rep 25: 2431-2446.e2437.

Marchionni E, Meneret A, Keren B, Melki J, Denier C, Durr A, Apartis E, Boespflug-Tanguy O, Mochel F. 2019. KIF1C Variants Are Associated with Hypomyelination, Ataxia, Tremor, and Dystonia in Fraternal Twins. Tremor Other Hyperkinet Mov (N Y) 9.

Merz C, Urlaub H, Will CL, Lührmann R. 2007. Protein composition of human mRNPs spliced in vitro and differential requirements for mRNP protein recruitment. RNA 13: 116–128.

Messitt TJ, Gagnon JA, Kreiling JA, Pratt CA, Yoon YJ, Mowry KL. 2008. Multiple kinesin motors coordinate cytoplasmic RNA transport on a subpopulation of microtubules in Xenopus oocytes. Dev Cell 15: 426–436.

Olsen JV, de Godoy LM, Li G, Macek B, Mortensen P, Pesch R, Makarov A, Lange O, Horning S, Mann M. 2005. Parts per million mass accuracy on an Orbitrap mass spectrometer via lock mass injection into a C-trap. Mol Cell Proteomics 4: 2010–2021.

Perry RB, Fainzilber M. 2014. Local translation in neuronal processes--in vivo tests of a “heretical hypothesis”. Dev Neurobiol 74: 210–217.

Pichon X, Moissoglu K, Coleno E, Wang T, Imbert A, Robert MC, Peter M, Chouaib R, Walter T, Mueller F et al. 2021. The kinesin KIF1C transports APC-dependent mRNAs to cell protrusions. RNA 27: 1528–1544.

Santos M, Damasio J, Carmona S, Neto JL, Dehghani N, Guedes LC, Barbot C, Barros J, Bras J, Sequeiros J et al. 2022. Molecular Characterization of Portuguese Patients with Hereditary Cerebellar Ataxia. Cells 11.

Schlager MA, Kapitein LC, Grigoriev I, Burzynski GM, Wulf PS, Keijzer N, de Graaff E, Fukuda M, Shepherd IT, Akhmanova A et al. 2010. Pericentrosomal targeting of Rab6 secretory vesicles by Bicaudal-D-related protein 1 (BICDR-1) regulates neuritogenesis. EMBO J 29: 1637–1651.

Schlautmann LP, Gehring NH. 2020. A Day in the Life of the Exon Junction Complex. Biomolecules 10.

Simpson JC, Joggerst B, Laketa V, Verissimo F, Cetin C, Erfle H, Bexiga MG, Singan VR, Heriche JK, Neumann B et al. 2012. Genome-wide RNAi screening identifies human proteins with a regulatory function in the early secretory pathway. Nat Cell Biol 14: 764–774.

Singh G, Kucukural A, Cenik C, Leszyk JD, Shaffer SA, Weng Z, Moore MJ. 2012. The Cellular EJC Interactome Reveals Higher-Order mRNP Structure and an EJC-SR Protein Nexus. Cell 151: 915–916.

Song T, Zheng Y, Wang Y, Katz Z, Liu X, Chen S, Singer RH, Gu W. 2015. Specific interaction of KIF11 with ZBP1 regulates the transport of beta-actin mRNA and cell motility. J Cell Sci 128: 1001–1010.

St Johnston D. 2005. Moving messages: the intracellular localization of mRNAs. Nat Rev Mol Cell Biol 6: 363–375.

Sutton MA, Schuman EM. 2006. Dendritic protein synthesis, synaptic plasticity, and memory. Cell 127: 49–58.

Suzuki K, Bose P, Leong-Quong RY, Fujita DJ, Riabowol K. 2010. REAP: A two minute cell fractionation method. BMC Res Notes 3: 294.

Tange T, Shibuya T, Jurica MS, Moore MJ. 2005. Biochemical analysis of the EJC reveals two new factors and a stable tetrameric protein core. RNA 11: 1869–1883.

Theisen U, Straube E, Straube A. 2012. Directional persistence of migrating cells requires Kif1C-mediated stabilization of trailing adhesions. Dev Cell 23: 1153–1166.

Tom Dieck S, Hanus C, Schuman EM. 2014. SnapShot: local protein translation in dendrites. Neuron 81: 958-958.e951.

Tyanova S, Temu T, Sinitcyn P, Carlson A, Hein MY, Geiger T, Mann M, Cox J. 2016. The Perseus computational platform for comprehensive analysis of (prote)omics data. Nat Methods 13: 731–740.

Wang DO, Martin KC, Zukin RS. 2010. Spatially restricting gene expression by local translation at synapses. Trends Neurosci 33: 173–182.

Wang DO, Ninomiya K, Mori C, Koyama A, Haan M, Kitabatake M, Hagiwara M, Chida K, Takahashi SI, Ohno M et al. 2017. Transport Granules Bound with Nuclear Cap Binding Protein and Exon Junction Complex Are Associated with Microtubules and Spatially Separated from eIF4E Granules and P Bodies in Human Neuronal Processes. Front Mol Biosci 4: 93.

Wang J, Huo K, Ma L, Tang L, Li D, Huang X, Yuan Y, Li C, Wang W, Guan W et al. 2011. Toward an understanding of the protein interaction network of the human liver. Mol Syst Biol 7: 536.

Wang Z, Ballut L, Barbosa I, Le Hir H. 2018. Exon Junction Complexes can have distinct functional flavours to regulate specific splicing events. Sci Rep 8: 9509.

Woodward LA, Mabin JW, Gangras P, Singh G. 2017. The exon junction complex: a lifelong guardian of mRNA fate. Wiley Interdiscip Rev RNA 8.

Xing L, Bassell GJ. 2013. mRNA localization: an orchestration of assembly, traffic and synthesis. Traffic 14: 2–14.

Yucel-Yilmaz D, Yucesan E, Yalnizoglu D, Oguz KK, Sagiroglu MS, Ozbek U, Serdaroglu E, Bilgic B, Erdem S, Iseri SAU et al. 2018. Clinical phenotype of hereditary spastic paraplegia due to KIF1C gene mutations across life span. Brain Dev 40: 458–464.

