## Supplementary Data for "The kinesin motor KIF1C is a putative transporter of the exon junction complex in neuronal cells"

Supplementary Figures:

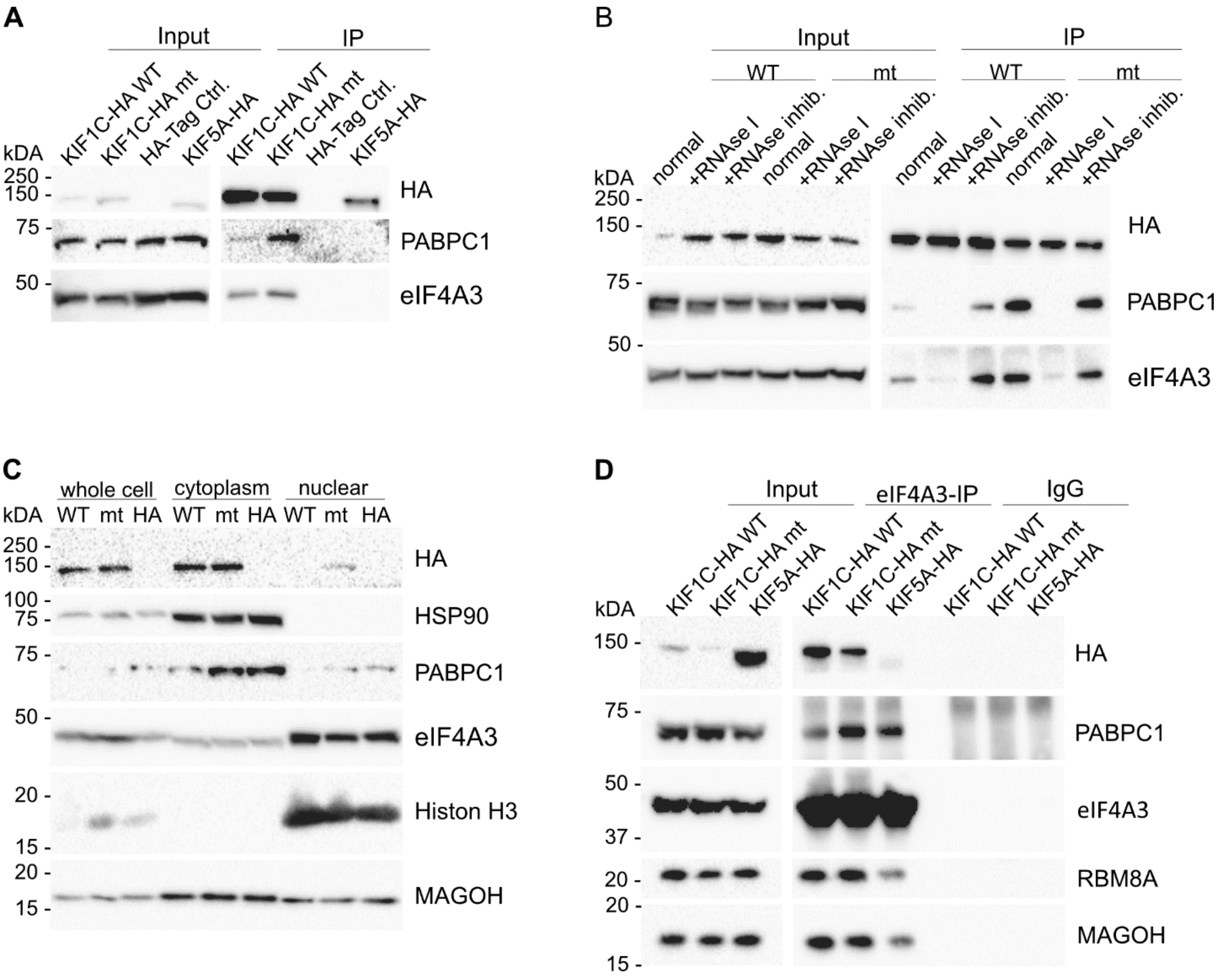


**Supplementary Figure S1: Supplementary HEK 293 cell data. A:** Immunoprecipitation of KIF1C complexes in HEK 293 cells stably overexpressing KIF1C-HA. PABPC1 and eIF4A3 were detected in the wildtype and mutant KIF1C IPs and were absent in the HA-Tag and KIF5A-HA control. **B:** RNAse IP was performed in HEK 293 cells overexpressing HA tagged $KIF1C_{WT}$ or $KIF1C_{G102A}$. The proteins interacted in a normal IP with KIF1C variants and the interaction was lost when the samples were treated with RNAse I before the IP, whereas a RNAse inhibition could restore the interaction. **C:** Fractionation was performed in HEK 293 cells overexpressing HA tagged $KIF1C_{WT}$ or $KIF1C_{G102A}$. As controls, Histon H3 was detected as a marker for the nuclear fraction and HSP90 for the cytoplasmic fraction. The HA tagged KIF1C variants as well as eIF4A3 were detected in all fractions. For PABPC1 the strongest signal was detected in the cytoplasmic fraction. **D:** Co- Immunoprecipitation of eIF4A3 in stable HEK 293 cell lines. $KIF1C_{WT}$ and $KIF1C_{G102A}$ were detected in the Co-IP and KIF5A-HA (control) was missing. All other tested proteins interacted with eIF4A3 as well and were therefore found in all IP samples.

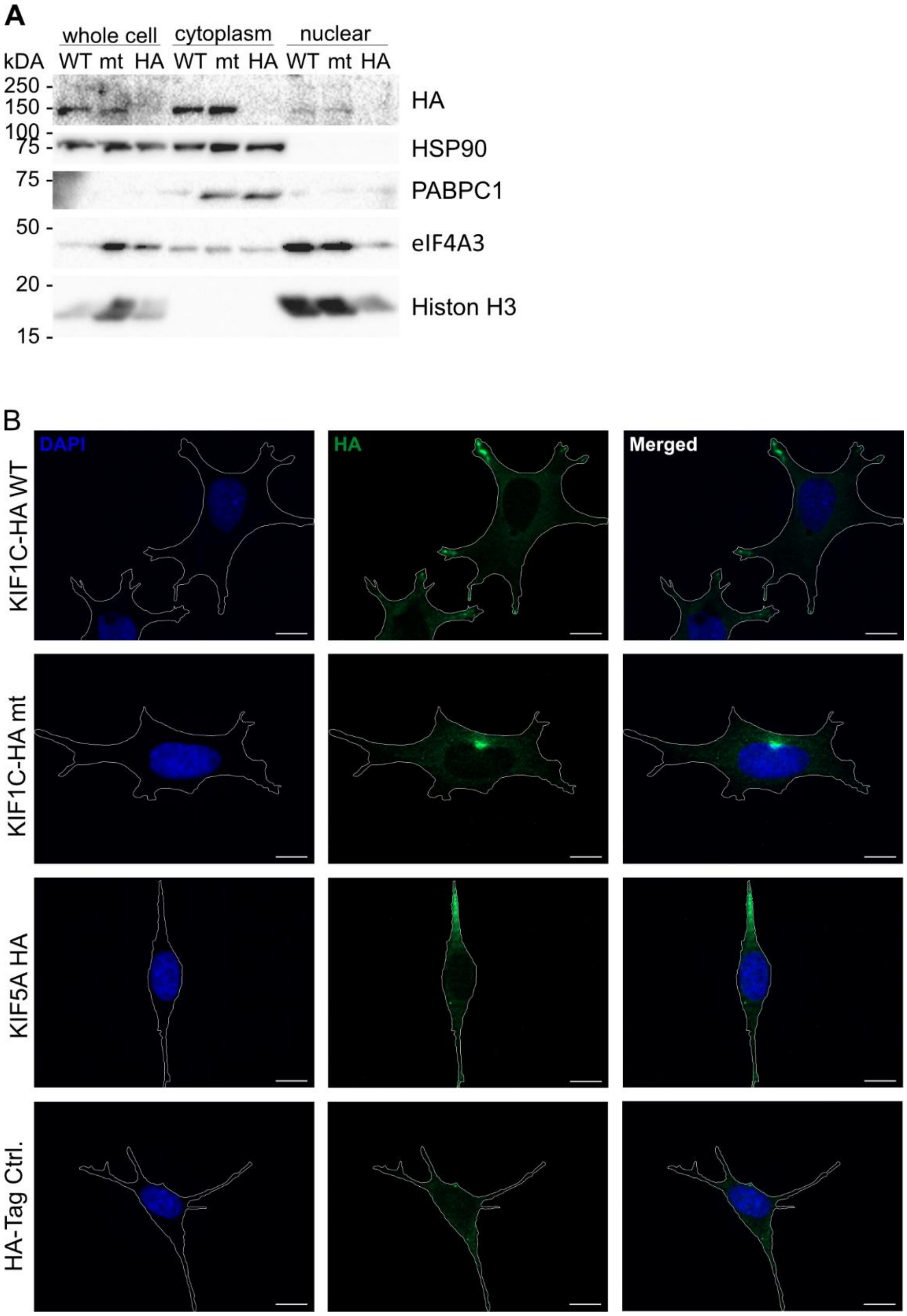


**Supplementary Figure S2: Supplementary SH-SY5Y cell data. A:** Fractionation was performed in differentiated SH-SY5Y cells overexpressing HA tagged KIF1C$_{WT}$ or KIF1C$_{G102A}$. As controls, Histon H3 was detected for the nuclear fraction and HSP90 for the cytoplasmic fraction. The HA tagged KIF1C variants were detected in all fractions, as well as eIF4A3. For PABPC1 the strongest signal was detected in the cytoplasmic fraction. **B:** Immunofluorescence was performed with all stable SH-SY5Y cell. The scale bars represent 10 µm and dashed lines indicate the outline of the cell body. The HA staining shows the localization of KIF1C$_{WT}$ in the tips of the cells, whereas KIF1C$_{G102A}$ localized in the pericentrosomal region. Wildtype KIF5A-HA also localizes at cell protrusions.

### Supplementary Table:

**Supplementary Table: MS Data**. The table shows all proteins identified in our MS analysis. The strict sig. hits (Significance A and permutation-based corrected t-test $p<0.05$) are marked in color. The hits only significant for $KIF1C_{WT}$ vs HA are marked in yellow, the hits significant for mutated $KIF1C_{G102A}$ vs HA are marked in blue and the overlay is marked in green.
